# Homeostatic activity-dependent tuning of recurrent networks for robust propagation of activity

**DOI:** 10.1101/033548

**Authors:** Julijana Gjorgjieva, Jan Felix Evers, Stephen J. Eglen

## Abstract

Developing neuronal networks display spontaneous rhythmic bursts of action potentials that are necessary for circuit organization and tuning. While spontaneous activity has been shown to instruct map formation in sensory circuits, it is unknown whether it plays a role in the organization of motor networks that produce rhythmic output. Using computational modeling we investigate how recurrent networks of excitatory and inhibitory neuronal populations assemble to produce robust patterns of unidirectional and precisely-timed propagating activity during organism locomotion. One example is provided by the motor network in *Drosophila* larvae, which generates propagating peristaltic waves of muscle contractions during crawling. We examine two activity-dependent models which tune weak network connectivity based on spontaneous activity patterns: a Hebbian model, where coincident activity in neighboring populations strengthens connections between them; and a homeostatic model, where connections are homeostatically regulated to maintain a constant level of excitatory activity based on spontaneous input. The homeostatic model tunes network connectivity to generate robust activity patterns with the appropriate timing relationships between neighboring populations. These timing relationships can be modulated by the properties of spontaneous activity suggesting its instructive role for generating functional variability in network output. In contrast, the Hebbian model fails to produce the tight timing relationships between neighboring populations required for unidirectional activity propagation, even when additional assumptions are imposed to constrain synaptic growth. These results argue that homeostatic mechanisms are more likely than Hebbian mechanisms to tune weak connectivity based on local activity patterns in a recurrent network for rhythm generation and propagation.

## Introduction

Activity-dependent adjustment of nascent synaptic connectivity is a widespread mechanism for tuning network properties in various parts of the developing brain (Kirkby et al., 2013). This activity is relayed by ordered topographical maps to higher processing centers, where it triggers correlated synaptic release from neighboring axonal terminals (Feller, 2009; Huberman et al., 2008; Katz and Shatz, 1996; Tritsch et al., 2007). This timing information is evaluated by postsynaptic neurons to sharpen their connectivity profile through Hebbian plasticity and thus improves network function during this early phase of circuit development.

Locomotor circuits also generate spontaneous activity during embryogenesis in many organisms, including chick, mouse, fish and fly (Crisp et al., 2008; Hanson and Landmesser, 2004; Kirkby et al., 2013; O’Donovan, 1999). These circuits are not topographically organized, yet endogenous activity patterns and synaptic release are required for coordinated motor behavior to develop (Crisp et al., 2011; Giachello and Baines, 2015; Warp et al., 2012). In *Drosophila*, motor neurons receive variable amounts of presynaptic input, suggesting that connectivity is plastic and acquires functional patterns in a homeostatically-regulated manner (Couton et al., 2015; Tripodi et al., 2008). The underlying mechanisms that might evaluate this early spontaneous activity and establish neuronal connectivity appropriately are unclear.

Computational modeling offers an efficient approach to explore the nature of the mechanisms that could underlie activity-dependent tuning of connectivity in a generic motor network that achieves a particular function. Here we study the organization of one such motor network that generates rhythmic behavior, the *Drosophila* larvae. Larvae crawl by peristaltic waves of muscle contractions which propagate along their body axis (Berni et al., 2012; Heckscher et al., 2012). These coordinated waves travel from posterior to anterior during forward locomotion, and from anterior to posterior during backward locomotion. The locomotor behavior is produced by central pattern generators (CPGs), which are segmentally repeated and modulated by sensory feedback, but can produce coordinated output even when sensory neurons are removed (Crisp et al., 2008; Fox et al., 2006; Suster and Bate, 2002). Mature crawling behavior of *Drosophila* larvae is preceded by uncoordinated yet neurally-controlled muscle contractions and incomplete peristaltic waves during late embryogenesis, which gradually improve before hatching (Crisp et al., 2008; Suster and Bate, 2002). Manipulating endogenous activity during this period can significantly affect the output of the circuit, suggesting that this activity helps to refine connectivity (Crisp et al., 2011; Giachello and Baines, 2015).

We previously established a model for wave propagation which produces propagating activity patterns with appropriate timing as during crawling in *Drosophila* larvae (Gjorgjieva et al., 2013). Although distinct connectivity parameters were shown to produce propagating waves of activity, the solution regions are complex (Figure 5 in Gjorgjieva et al. (2013)), and it is unclear how they arise during development. Here, we examine two different activity-dependent mechanisms which can tune network connectivity during development: (1) a Hebbian plasticity model, where coincident activity between neighboring neuronal populations strengthens connections between them, and (2) a homeostatic model, where synaptic connections are modified to maintain a constant level of excitatory postsynaptic activity based on spontaneous input. We show that homeostatic mechanisms are more appropriate than Hebbian mechanisms to organize connectivity in these motor networks. We demonstrate the case by comparing peristaltic wave properties for the two models based on the final configuration of network connectivity produced by the models. We also demonstrate the robustness of activity-dependent tuning of connectivity by varying the properties of spontaneous activity to predict how manipulations of this activity during embryogenesis might impact output of the mature network. Thus, our work highlights the relative importance of homeostatic mechanisms for establishing functional connectivity in developing motor networks, in contrast to sensory networks where connectivity is more tightly constrained enabling the formation of accurate sensory maps.

## Materials and Methods

### Network model

The network model was based on the interaction of recurrently coupled excitatory and inhibitory populations to produce robust propagation of activity with appropriate timing relationships as observed experimentally during crawling in *Drosophila* larvae (Gjorgjieva et al., 2013). Such unidirectional propagation of activity is also observed in other experimental systems (see Discussion). Our model is an abstraction of neural circuits that might generate larval crawling and captures the observed coordination at the segmental level. The model contains eight coupled units of excitatory (*E*) and inhibitory (*I*) subpopulations to represent the activity in the eight abdominal segments of the larva (Figure 1A).

**Figure 1.**
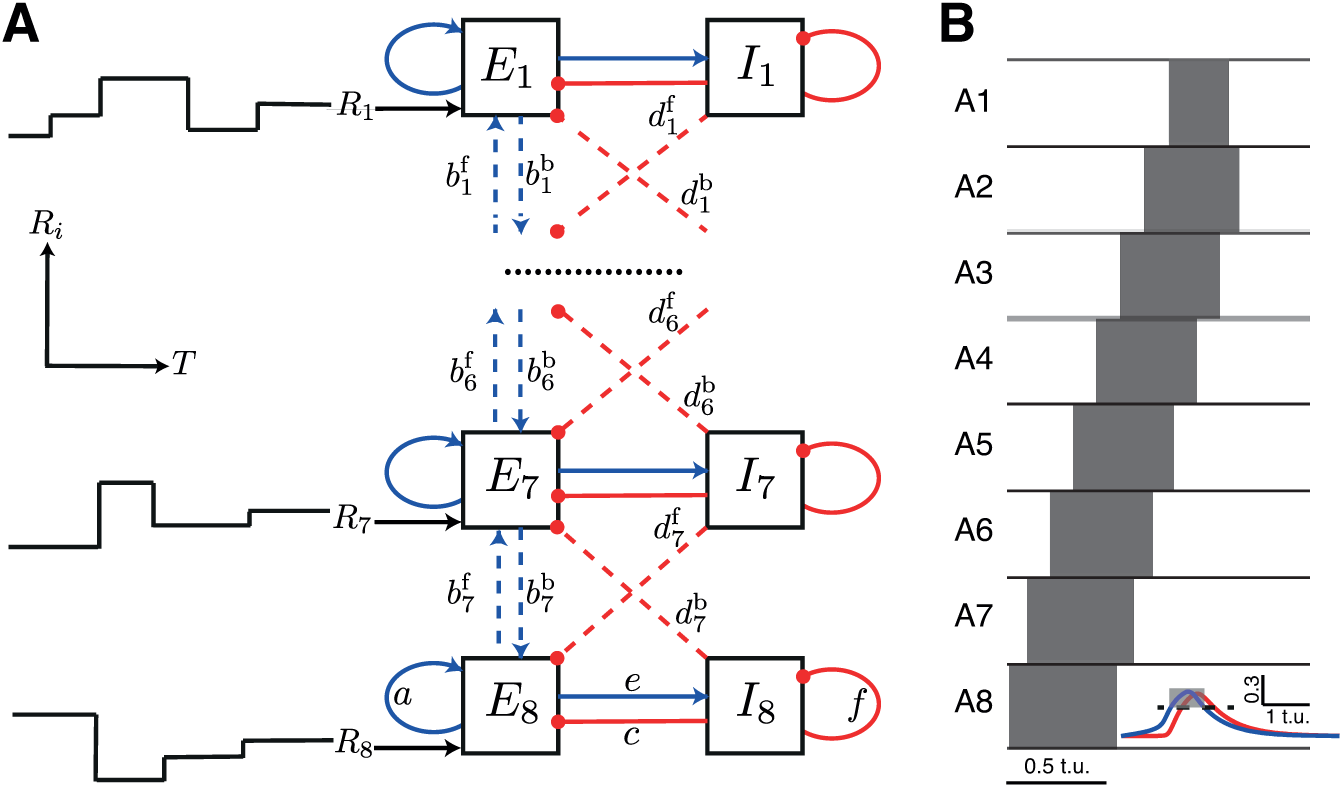
A network model for the emergence of coordinated output. **A**. A network model for peristaltic wave propagation with eight segments, each consisting of an excitatory (*E*) and an inhibitory (*I*) neuronal population. The connections within the same segment (solid lines), *a*, *c*, *e* and *f*, shown only for the most posterior segment, are fixed during development. Nearest-neighbor segments are connected with excitatory, *b* (blue, arrows), and inhibitory, *d* (red, dots), connections. These connections are plastic during development as indicated by the dashed lines. The weights for forward and backward wave propagation are identified by superscripts ‘f’ and ‘b’, respectively. Each segment receives spontaneous random input of strength *R_i_* for a duration of *T_i_*, as illustrated on the left. **B**. An example of supra-threshold excitatory activity (threshold for detection of supra-threshold activity is 0.3) of each segment. Time is measured in arbitrary time units (t.u.). The inset shows excitatory activity (blue), and inhibitory activity (red) for segment A8 with a threshold denoted by the dashed line.

The activity in each segment was modeled with a Wilson-Cowan unit (Wilson and Cowan, 1972) consisting of two neuronal populations, excitatory (*E*) and inhibitory (*I*). These two populations represent the joint activity of all central neurons in the CPG circuit for crawling, but are sufficiently general to represent excitatory and inhibitory neurons in other systems. The differential equations for the time-dependent variation of averaged excitatory and inhibitory neuronal activities were

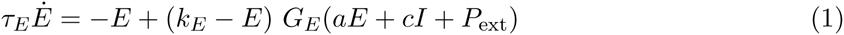

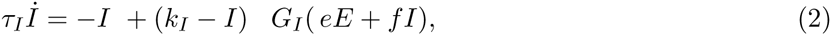
where the functions *G_E_*, *G*_1_ and *G_S_* represent sigmoidal response functions of the excitatory, inhibitory and sensory neuronal populations given by

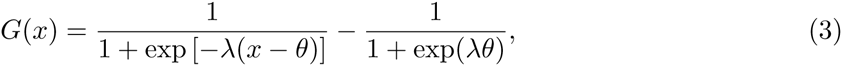
λ represents the maximum slope of the sigmoid (or if *G* represents an activation function, it denotes the speed of activation) and *θ* represents the location of the maximum slope (or the threshold for activation). The terms *k_E_* and *k_I_* denote the maxima of the response functions for the excitatory and inhibitory populations, *k_E_* = 0.9945 and *k_I_* = 0.9994. The time constants *τ_E_* and *τ_I_* control decay of excitatory and inhibitory activities after stimulation, and determine the timescale of network activity. The connectivity coefficients *a* (or *e*) and *c* (or *f*) represent the average number of excitatory and inhibitory synapses per cell in the excitatory (or inhibitory) population, respectively. The time-varying function *P*_ext_ (*t*) denotes the external input applied to the excitatory population of one of the end segments to initiate a wave. Note that this input was not used during development in the Hebbian and homeostatic models; it was only applied to test whether a network was functional at the end of development.

To model wave propagation, we coupled eight Wilson-Cowan *E-I* units. Two types of connections were created between neighboring segments. First, bidirectional excitatory connections (*b*) between excitatory populations of neighboring segments allow activity to propagate along segments. Second, inhibitory connections (*d*) from the inhibitory population in one segment to the neighboring (anterior and posterior) excitatory populations terminate activity in each previously active segment and ensure unidirectional wave propagation (Figure 1). This network was equipped with symmetric bidirectional connectivity such that forward and backward propagating waves were generated with similar properties, consistent with experiments (Gjorgjieva et al., 2013). We list parameter values that generate a functional wave (assuming appropriate connectivity) in Table 1.

**Table 1.** Network parameters for wave propagation. Default parameter values for the model simulations.

| $a$ | $b$ | $c$ | $d$ | $e$ | $f$ | $\tau_E$ | $\tau_I$ | $P_{\text{ext}}$ | $b_E$ | $b_I$ | $\theta_E$ | $\theta_I$ |
| --- | --- | --- | --- | --- | --- | --- | --- | --- | --- | --- | --- | --- |
| 16 | 20 | -12 | -20 | 15 | -3 | 0.5 | 0.5 | 1.7 | 1.3 | 2 | 4 | 3.7 |

A ‘contraction’ in the model was defined as supra-threshold activity (activity above a threshold, 0.2 or 0.3) of the excitatory population (Figure 1B, horizontal dashed line). Such contractions were used to implement the activity-dependent weight refinement in the Hebbian and homeostatic development models. We examined the timing relations between neighboring segments during propagating waves of supra-threshold activity in the model by analyzing two quantities: The *duty cycle* was the total time that each excitatory activity is supra-threshold relative to the wave duration. The *interburst interval* was the time from the initiation of supra-threshold activity of one segment to the initiation of supra-threshold activity of the neighboring segment, normalized by wave duration.

Weights are labeled with a superscript ‘f’ or ‘b’, to denote their respective role in forward or backward wave propagation (Figure 1). Thus, there were 28 modifiable weights:

- seven 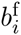 weights connecting *E_i_*_+1_ to *E_i_*,
- seven 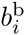 weights connecting *E_i_* to *E_i_*_+1_,
- seven 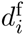 weights connecting *I_i_* to *E_i_*_+1_, and
- seven 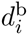 weights connecting *I_i_*_+1_ to *E_i_*, where *i* = {1,… 7}.

The model was initialized with weak connectivity incapable of generating waves. We considered two scenarios: (1) All the weights started identically *b* = −*d* = 2; this was 10% of the values used to generate robust waves (Gjorgjieva et al., 2013); (2) The weights were independent random samples drawn from a uniform distribution in the range between 0 and 5 for excitatory weights, and −5 to 0 for inhibitory weights. The final value of the weights was somewhat dependent on the initial conditions, and in general, the steady state weights showed greater variability when starting from random initial conditions. However, the stability of the weights was unaffected. The other weights within the same segment, *a, c, e* and *f*, were kept fixed as in Table 1. All parameters were dimensionless and time in the model was measured in arbitrary time units (t.u.).

### Time of development

For both activity-dependent models simulations of development were conducted for a total time of 2 × 10^5^ t.u. The total simulation time depends on the time constant of weight update, the nature of spontaneous activity and how spontaneous activity changes during development. The notion of ‘developmental time’ in the model is useful in describing the gradual improvement of coordination in the activity patterns of the developing network.

### Generating spontaneous activity

As there is little quantitative characterization of spontaneous activity in the *Drosophila* motor network during development, here we made simple approximations to generate spontaneous activity. Inputs of strength *R_i_* for duration *T_i_* were applied to each excitatory population *E_i_* (Figure 1). When the spontaneous input into a given population *E_i_* was sufficiently strong, *E_i_* became supra-threshold. At the time of threshold-crossing, activity-dependent mechanisms were applied to modify the weights. The duration of each spontaneous input, *T_i_*, was a random number drawn from a uniform distribution, *U*(*x*_1_, *x*_2_), determining the total time for which the input of strength *R_i_* was applied to *E_i_* before choosing a new spontaneous input. Sampling the duration of spontaneous input from a distribution ensured that neuronal activity in different populations crossed threshold at different times as we did not want to introduce correlations in spontaneous activity. Since network connectivity is recurrent, the network itself generates a significant proportion of activity when the weights are sufficiently strong. This automatically generates correlations between the activations of neighboring segments, even though spontaneous activity itself is uncorrelated. The strength of spontaneous input, *R_i_*, was also a random number; this input was applied to *E_i_* for time *T_i_*. We used a truncated Gaussian distribution with mean *μ* and variance *σ*^2^, [*N*(*μ*, *σ*)]+, to ensure that spontaneous input remained positive.

The parameters of the distribution for *T_i_* did not greatly affect the frequency of supra-threshold events in a given excitatory population (data not shown). In contrast, *R_i_* had a strong effect upon the frequency of threshold crossings in the model network. This was partially because weights were modified precisely at the time when population activity crossed threshold. Therefore, we explored the effect of the distribution of *R_i_* on the outcome of weight development in the two plasticity models.

During development of *Drosophila* embryos, frequent random activity uncoordinated across segments gradually becomes replaced by coordinated activity ranging across several neighboring segments, with all contraction activity ceasing shortly before hatching when the larva emerges from the egg shell and starts to crawl (Crisp et al., 2008). This is consistent with a model where spontaneous activity decreases during development. To model this effect, we considered decreasing either the mean or the variance of the Gaussian distribution for *R_i_*. In the Hebbian model we decreased the standard deviation in steps; however, the model was insensitive to how spontaneous activity was reduced. The homeostatic model was also robust to the decrease of spontaneous activity, though the speed of decrease of spontaneous activity affected the speed at which weights stabilized.

### The Hebbian bidirectional model

Weight updates followed the rate-based formulation of Hebbian plasticity: if presynaptic precedes postsynaptic activity then the corresponding weight is potentiated, else the weight is depressed. For any two populations connected by a weight the activity which crossed threshold first was presynaptic. Weights were modified only for the direction in which a wave would have propagated had the connection between the two populations been sufficiently strong (Figure 2A,B). Namely, if *E_i_*_+1_ crossed threshold before *E_i_*, then 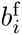 was potentiated, while 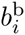 was depressed. To implement potentiation, a small constant, Δ_+_ > 0, was added to the weight

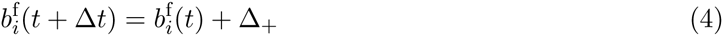
and similarly, if the constant for weight depression was denoted by Δ_−_ > 0, 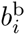depressed.

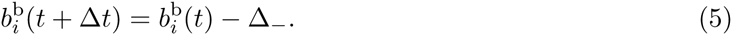

**Figure 2.**
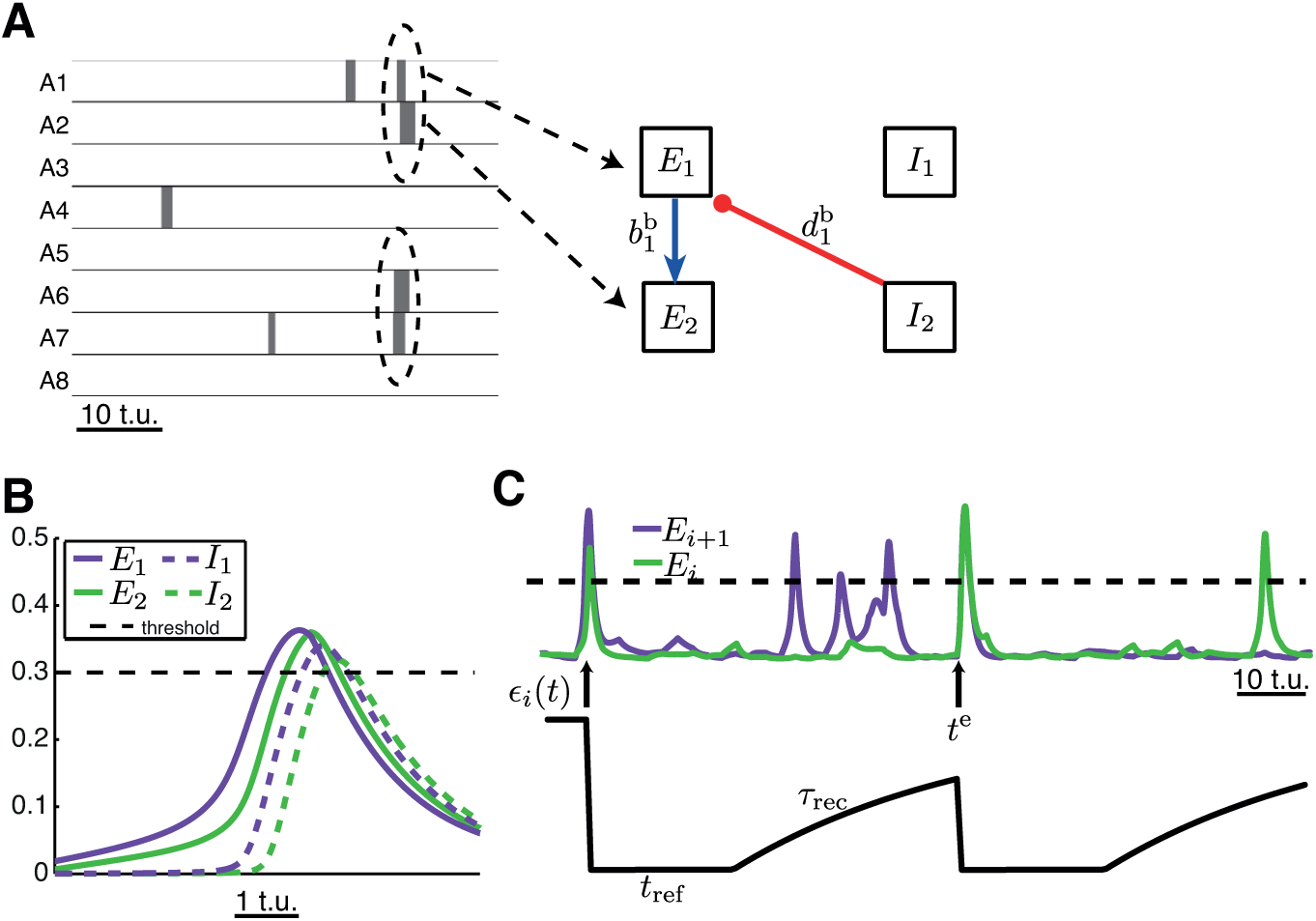
The Hebbian model for synaptic modification with synaptic efficacy. **A**. Weight changes induced by Hebbian plasticity: A snapshot of 50 time units of activity during early development. When activity in neighboring neuronal populations exceeds threshold simultaneously (dashed ellipses), based on the order of threshold crossing, the respective forward or backward weights are updated, here 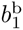 and 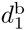. **B**. Excitatory and inhibitory activity in two neighboring segments simultaneously above threshold is shown as a function of time. **C**. As the activity of two neuronal populations crossed threshold (dashed line) simultaneously, at times denoted by *t^e^* (arrows), the efficacy *∊_i_*(*t*) was reset to 0 for a refractory period of *t*_ref_ and then exponentially recovered to 1 with a timescale *τ*_rec_.

If *E_i_* crossed threshold before *E_i_*_+1_, then 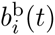 potentiated

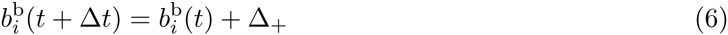
and 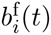 depressed

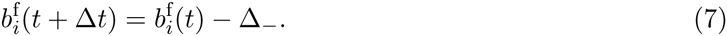

For the inhibitory weights, if *I_i_* crossed threshold before *E_i_*_+1_, then 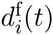 strengthened

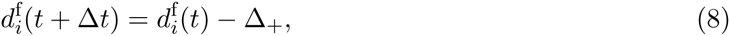
while if *E_i_*_+1_ crossed threshold before *I_i_*, then 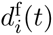 weakened

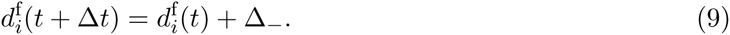

And similarly if *I_i_*_+1_ crossed threshold before *E_i_*, then 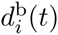 strengthened

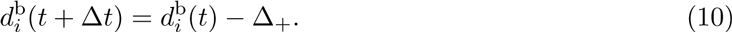
while if *E_i_* crossed threshold before *I_i_*_+i_, then 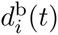 weakened

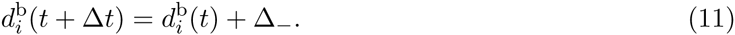

For a weight to be modified, activity of the pre- and postsynaptic populations connected by the weight had to be simultaneously supra-threshold. Weight changes were applied instantaneously, immediately after the second of the two populations connected by the weight crossed threshold. However, the model was robust to this requirement, and the weight could also be updated as the activity of each population dropped below threshold.

### The Hebbian efficacy model

In the Hebbian efficacy model, synchronous threshold crossing of activity in neighboring populations resulted only in weight potentiation and no depression. To ensure that weights did not grow without a bound, weight increase was limited by the frequency of contraction of the neuronal populations connected by the synaptic weight. Such activity-dependent control of the weights was motivated by the regulation of activity-dependent depression in developing motor networks (Crisp et al., 2008; Fedirchuk et al., 1999).

To implement this activity-dependent synaptic depression, a ‘synaptic efficacy’ variable was introduced in the Hebbian model (Froemke and Dan, 2002). This efficacy variable ∊ depressed to 0 at the time when the activity of a neuronal population crossed threshold (Figure 2C). Then, in equations (4) and (6), when the excitatory activity in two populations crossed threshold, the excitatory weights, 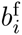 and 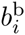, were respectively modified by 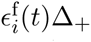 and 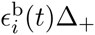. Similarly, instead of Eqs. (8) and (10), the inhibitory weights, 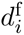 and 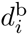, were decreased by 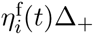 and 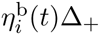, respectively, with efficacies

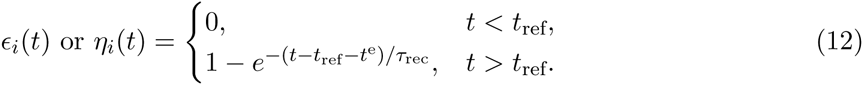

Here, *t^e^* denotes the time when activity of the two neuronal populations connected by the weight crossed threshold (Figure 2C). The efficacy was held at 0 for a refractory time period *τ*_ref_, after which it recovered to 1 with a time constant of *τ*_rec_. The refractory period was implemented to ensure that after the weights had strengthened stufficiently to generate waves, threshold crossings of the activity in neuronal populations during propagating waves did not further modify the weights. The value of *τ*_rec_ was chosen such that the period between two consecutive waves was larger than the refractory period *τ*_ref_.

To equalize the frequency of threshold crossing among the different excitatory populations, an additional ‘end drive’ was applied to the end excitatory populations, *E*_8_ and *E*_1_ (Figure 5). This drive was applied to one of *E*_8_ or *E*_1_ chosen randomly for a duration *T_i_* = 3 t.u. instead of the spontaneous input *R_i_*. We assumed the strength of this end drive increased as spontaneous activity decreased during development, reaching a final value of 1.7.

### The homeostatic model

The homeostatic model was implemented as in the Bienenstock-Cooper-Munroe (BCM) theory for synaptic plasticity (Bienenstock et al., 1982). Weight modification was bidirectional (potentiation or depression) to maintain a constant averaged activity of each excitatory population, *r_i_*

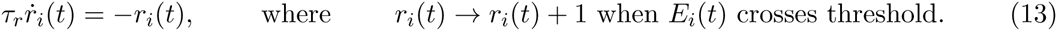

The time constant *τ_r_* was chosen to be much longer than the timescale of neuronal dynamics. Weight potentiation or depression depended on whether the average excitatory activity in a given segment was greater or smaller than a threshold for synaptic modification, *θ*. The homeostatic model is activity-dependent, thus in the absence of spontaneous activity (*μ* → 0) the weights in the network will not be modified. The threshold is defined to be a function of the averaged excitatory activity itself

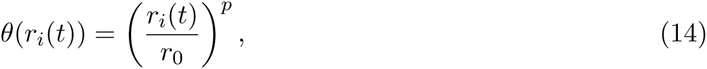
where *r*_0_ = 2.0 was a constant and *p* > 1 (here, *p* = 1.5). The modification function *ϕ* depends on *r_i_*(*t*) and *θ_i_*(*r_i_*(*t*))

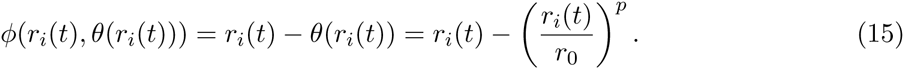

Then, synaptic weight change was implemented according to

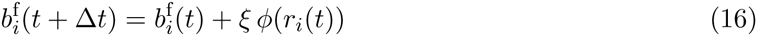
for the forward excitatory weights, and

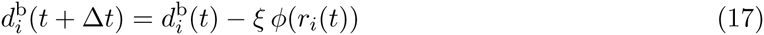
for the backward inhibitory weights for which *E_i_* is postsynaptic (*i* = {1, 2,…, 7}). Similarly,

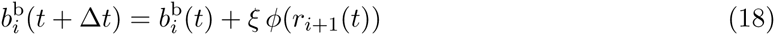
for the backward excitatory weights, and

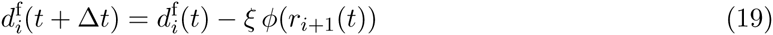
for the forward inhibitory weights for which *E_i_*_+1_ is postsynaptic (*i* = {1, 2,…, 7}). The constant *ξ* was a small positive number to ensure weights change slowly relative to the timescale of neuronal dynamics (e.g. 0.001). In the limit of *r_i_*(*t*) ≪ *r*_0_,

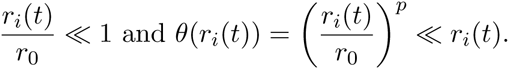

This leads to *ϕ*(*r_i_*(*t*), *θ*(*r_i_*(*t*))) > 0 since *p* > 1, and potentiation of the weights. In the limit of *r_i_*(*t*) ≫ *r*_0_,

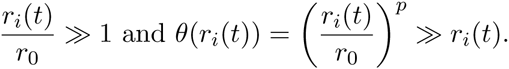

This leads to *ϕ*(*r_i_*(*t*), *θ*(*r_i_*(*t*))) < 0, and depression of the weights. In both models, the weights were prevented from changing sign in agreement with Dale’s law (Strata and Harvey, 1999).

## Results

Can spontaneous activity successfully tune a bidirectionally connected recurrent network with weak connectivity by using local activity-dependent mechanisms that modify connection strength to produce functional output? Here we studied the generation of unidirectional propagating waves of activity resembling those observed during crawling in the *Drosophila* larval motor network. Two main principles provided the basis for our developmental model: (1) The model should use local activity-dependent mechanisms to tune connection strength, and (2) The connections should achieve a stable configuration at the end of development such that the network generates propagating waves of activity with regular timing relationships (interburst intervals and duty cycles) in different segments.

We used a recurrent network with bidirectional connectivity which can generate forward and backward propagating waves when appropriately tuned (Figure 1, see also Figure 3 in Gjorgjieva et al. (2013)). The model was initialized with weak connectivity between different segments incapable of generating waves. Weak connectivity in the network is likely specified by activity-independent mechanisms, which guide axons toward their correct target partners. These guidance mechanisms are widespread throughout the central nervous system (Dickson and Gilestro, 2006; Hand and Kolodkin, 2015; Huberman et al., 2008; Klein and Kania, 2014). Each excitatory population in the model network was triggered by patterned spontaneous input (see Methods), and depending on its strength and duration, increased the activity of the populations in random segments above threshold at different times. We sought to determine the nature of activity-dependent plasticity rules that modify the strength of excitatory and inhibitory weights, *b* and *d*, connecting neighboring segments using the spatio-temporal activation patterns of the different segments. Having as a goal to robustly produce propagating waves of activity with precise timing relationships, we compared a Hebbian to an activity-dependent homeostatic model for connection tuning based on local activity patterns.

**Figure 3.**
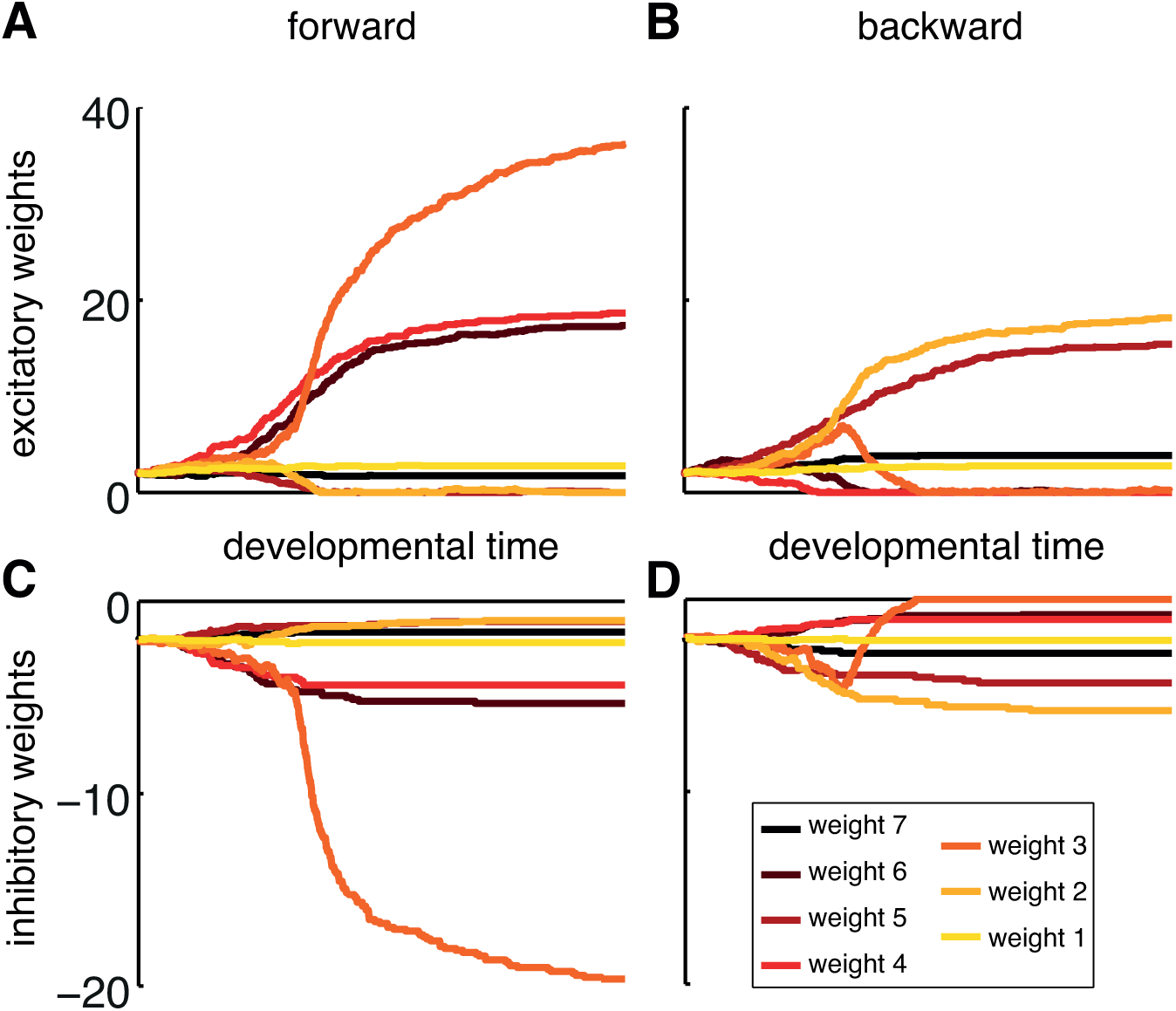
Weight development in the Hebbian bidirectional model without synaptic efficacy. Weights were modified using the Hebbian bidirectional model (Eqs. 4–11) with Δ_+_ = 0.1 and Δ_−_ = 0.05 without activity-dependent synaptic efficacy. Spontaneous activity was generated with *R_i_* ~ [*N*(0.2,0.8)] _+_ and *T_i_* ~ *U*(2, 3). Every 8,000 time units, the standard deviation (*σ* = 0.8) of the distribution for *R_i_* was decreased by 0.05. **A**. Forward excitatory weights 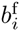, **B**. Backward excitatory weights 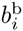, **C**. Forward inhibitory weights 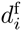 and **D**. Backward inhibitory weights 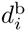.

### The Hebbian bidirectional model does not generate functional weight distributions

Weight modification in the Hebbian model was based on Hebb’s mechanism (Hebb, 1949): if the population activity in two neighboring segments is simultaneously above threshold, the weights between the neuronal populations should increase. Even though spontaneous activity in the model was uncorrelated across segments, correlations between the activations of neighboring segments exist because of the network’s recurrent connectivity. Hebb’s idea has been translated into various forms of correlation-based rules for synaptic modification, which commonly assume that synapses change strength in proportion to the correlation of the pre- and postsynaptic activity. This includes spike-timing-dependent plasticity (STDP), where the magnitude of synaptic potentiation and depression depends not only on the order of pre- and postsynaptic activity, but also on their timing (Markram et al., 1997). Hebbian plasticity has been successfully applied to activity-driven refinement of developing circuits and learning in neuronal networks in many brain regions, including hippocampus, neocortex, and cerebellum (Bliss and Collingridge, 1993; Bliss and Lomø, 1973; Malenka and Bear, 2004; Malenka and Nicoll, 2009; Martin et al., 2000). Correlation-based Hebbian rules have been often used to model map formation in the visual system (Miller, 1992; Miller et al., 1989). Therefore, we first characterize the role of Hebbian rules in connectivity tuning in the recurrent networks for motor output studied here.

A common challenge that Hebbian mechanisms face is unbounded growth of synaptic weights. Although upper bounds on synaptic strength can be imposed, and most biological systems probably have a saturation constraint limiting synaptic growth, it is unlikely that this upper bound is always fixed to the same value and that the system knows this value *a priori*. To prevent unbounded synaptic growth, we considered two implementations of the Hebbian model. One implementation is based on *bidirectional* plasticity, which evokes synaptic potentiation if presynaptic activity comes before postsynaptic activity, and depression if the order of activity is reversed (in line with STDP). This model has been referred to as the *Hebbian bidirectional model* (see Methods). The Hebbian bidirectional model fails to produce stable bidirectional connections because the weights develop in a biased way: some potentiate, while others decrease to zero, uncoupling neighboring populations (Figure 3).

What is the reason for this bias? Because random spontaneous input is applied to each excitatory population in the network with initially small *b* and *d*, some weights randomly potentiate more than others. However, each pair of neighboring excitatory populations is connected by weights for wave propagation in both directions, so whenever the weight for one direction potentiates, the corresponding weight in the other direction depresses. This creates a feedback loop, in which stronger weights produce more frequent supra-threshold activity, further amplifying the same sets of weights, while the weights in the opposite direction decay to zero (Figure 3). Therefore, it is not possible to strengthen synaptic connections in both directions between neighboring excitatory populations appropriately for wave generation, because of the asymmetric integration window of the Hebbian model. This is consistent with other models of STDP (Abbott and Nelson, 2000; Song and Abbott, 2001). Non-linear plasticity rules can generate functional bidirectional connectivity, but such models are based on detailed spiking and voltage trajectories of individual neurons (Clopath et al., 2010).

### The Hebbian efficacy model requires balancing of activity across the network

We therefore considered an alternate implementation of the Hebbian model, where synaptic potentiation is induced if presynaptic activity occurs before postsynaptic activity, augmented by activity-dependent synaptic efficacy to prevent unbounded weight increase, but without any synaptic depression. This model has been referred to as the *Hebbian efficacy model*.

The amplitude of weight modification in the Hebbian efficacy model is inversely proportional to the frequency of threshold crossings in segments connected by that weight (see Methods). The model produces bidirectional weights that become ordered in the direction of wave propagation (Figure 4A). Thus, of all excitatory weights for forward propagation, the most posterior 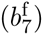 becomes the largest, while the weights in the middle of the network become the smallest (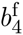 and 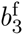). We found this to be due to a combination of (1) the difference in total input (synaptic and spontaneous) received by each excitatory population, and (2) the preference for synaptic potentiation of the forward weights induced by the order of excitatory segment activation – a hallmark of Hebb’s rule. Indeed, segment A8 has only one neighbor and receives overall less drive than the other segments in the network; thus, its efficacy is the largest, evoking the highest potentiation of 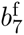. While the most anterior segment, A1, also has only one neighbor, it can also be driven from its neighboring segment (A2) in the forward direction so 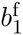 has a lower efficacy and does not potentiate as much. The weights in the middle are most frequently activated, they have the lowest efficacy and are potentiated the least.

**Figure 4.**
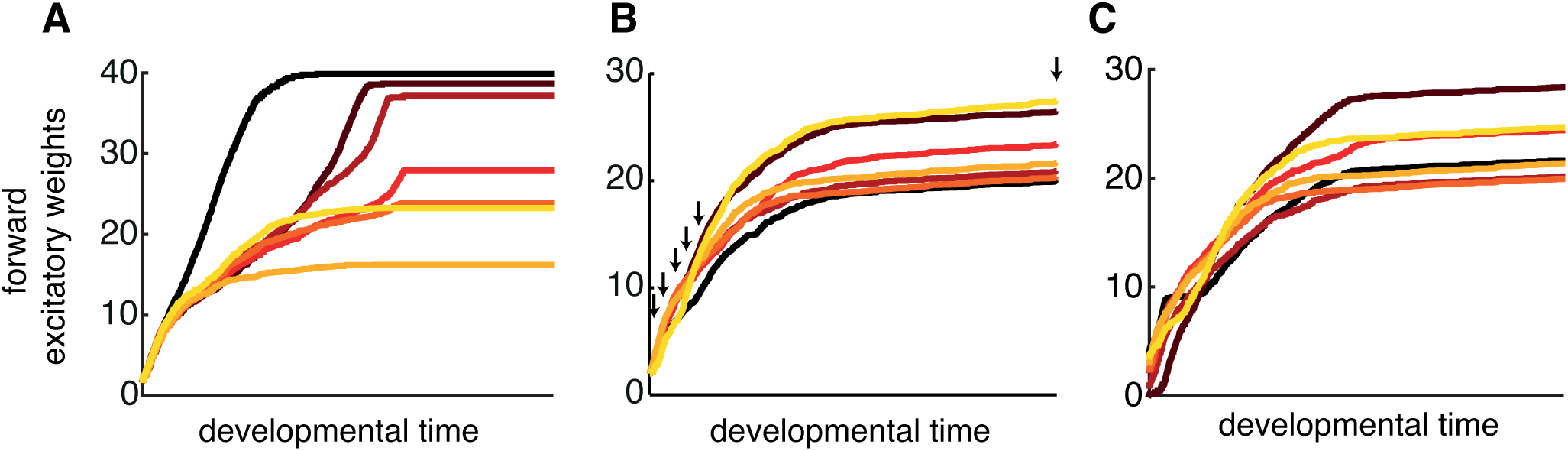
Weight development in the Hebbian efficacy model. **A**. The network weights were modified using the Hebbian efficacy model (Eq. 12), where *τ*_rec_ = 20 t.u. and *τ*_ref_ = 20 t.u. All weights start at the same initial condition (*b* = −*d* = 2). **B**. As A but with additional end drive applied to one of the end segments chosen randomly with equal probability (reaching 1.7 at the end of the simulation). Spatio-temporal patterns of excitatory activity at the times denoted with arrows are shown in Figure 6. **C**. Same as B but with random initial conditions starting in the range of 0 and 5. In all cases spontaneous activity was generated with *R_i_* ~ [*N*(0.5,0.8)]_+_ and *T_i_* ~ *U*(2, 3). Every 8,000 time units, the standard deviation (*σ* = 0.8) of the distribution for *R_i_* was decreased by 0.05. Only excitatory weights in the forward direction are shown as per Figure 3.

We examine this in greater detail. The threshold crossing frequency of excitatory activity (necessary to trigger weight change) depends strongly on the strength of the weights at a given time point during the development simulation (Figure 5A). When weights are small, threshold crossing is purely the result of spontaneous drive, which is approximately uniform across all segments – thus, all segments exhibit approximately the same number of threshold crossings (blue, Figure 5A). When the weights are sufficiently strong to generate waves, then the activation of any one segment propagates to all remaining segments, again activating all of them uniformly. Hence, all segments again exhibit approximately the same number of threshold crossings (red, Figure 5A). The biggest discrepancy between the frequency of supra-threshold activity in different segments occurs at intermediate synaptic strength (yellow, Figure 5A). In this scenario, the end segments of the network have only one neighboring segment and receive the lowest synaptic input, and so have the lowest overall input. Weights connecting neuronal populations whose activity crosses threshold more frequently due to the higher overall drive (in the middle of the network) have smaller efficacies compared to other weights which connect less active populations (at the ends of the network). Therefore, the former weights potentiate less than the latter. The outcome of this process produces variable weights (Figure 4A). Taking the weights at the end of the developmental period fails to produce propagating waves of activity across the network (data not shown).

**Figure 5.**
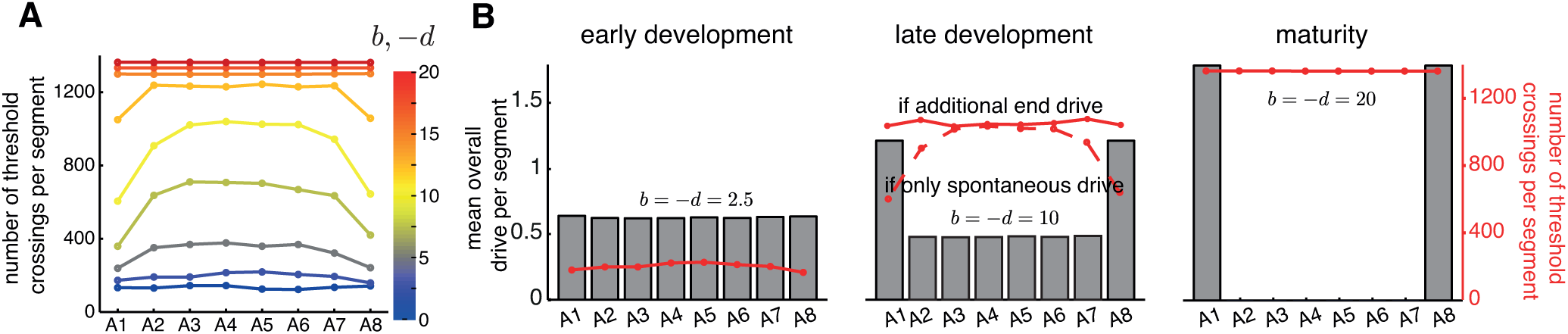
Frequency of threshold crossing of excitatory populations in the different models with Hebbian plasticity. **A**. Number of threshold crossings per segment for 10,000 time units where spontaneous activity was generated with *R_i_* ~ [*N*(0.2, 0.8)]+ and *T_i_* ~ *U*(0.5,1.5). Here spontaneous activity was fixed during the simulation. The strength of the intersegmental connections in the network (Figure 1) depends on the strength of the weights, which were taken to be uniform across the network. **B**. The total amount of activity received by each segment consists of different contributions from spontaneous input and from additional end drive at different points in development. The histograms show the mean overall drive per segment (spontaneous input and end drive) at three developmental time points (early development, late development and maturity) showing a decrease of spontaneous activity and an increase in end drive. The number of threshold crossings per segment is also shown (red) as in A for a simulation of 10,000 time units. The middle panel shows that with additional end drive (full line) all segments contract at a similar frequency, compared to the case with no end drive (dashed line).

If Hebbian-style activity-dependent mechanisms indeed tune synaptic connectivity in the bidirectionally connected network, then to equalize the total amount of input received by each segment, the end segments must receive additional spontaneous input (Figure 5B). Furthermore, as spontaneous activation of all segments decreases during development (Crisp et al., 2008), then the intensity of additional spontaneous drive to the end segments must increase. The increase in the end drive during development might correspond to the biological system progressively testing more frequently whether the network is fully assembled to produce waves. Thus, towards the of development, the only drive to the network is to the end segments which will initiate either forward or backward waves (Figure 5B).

### Weight distributions from the Hebbian efficacy model produce unreliable waves even with balanced activity

The Hebbian efficacy model with additional end drive successfully produces stable and functional weight distributions (Figure 4B). To examine the progression of network output during development, we recorded the spatio-temporal activity patterns in the network over a given period (Figure 6A–E) at the developmental time points denoted with arrows in Figure 4B. The network output gradually becomes more coordinated similarly to motor development in *Drosophila* (Crisp et al., 2008). Furthermore, partial motifs of coordinated output, which involve the simultaneous supra-threshold activity of excitatory populations in several neighboring segments, typically originate at one end of the network. This makes a concrete experimental prediction for how coordinated activity emerges gradually during development if the Hebbian model with additional end drive organizes the network.

**Figure 6.**
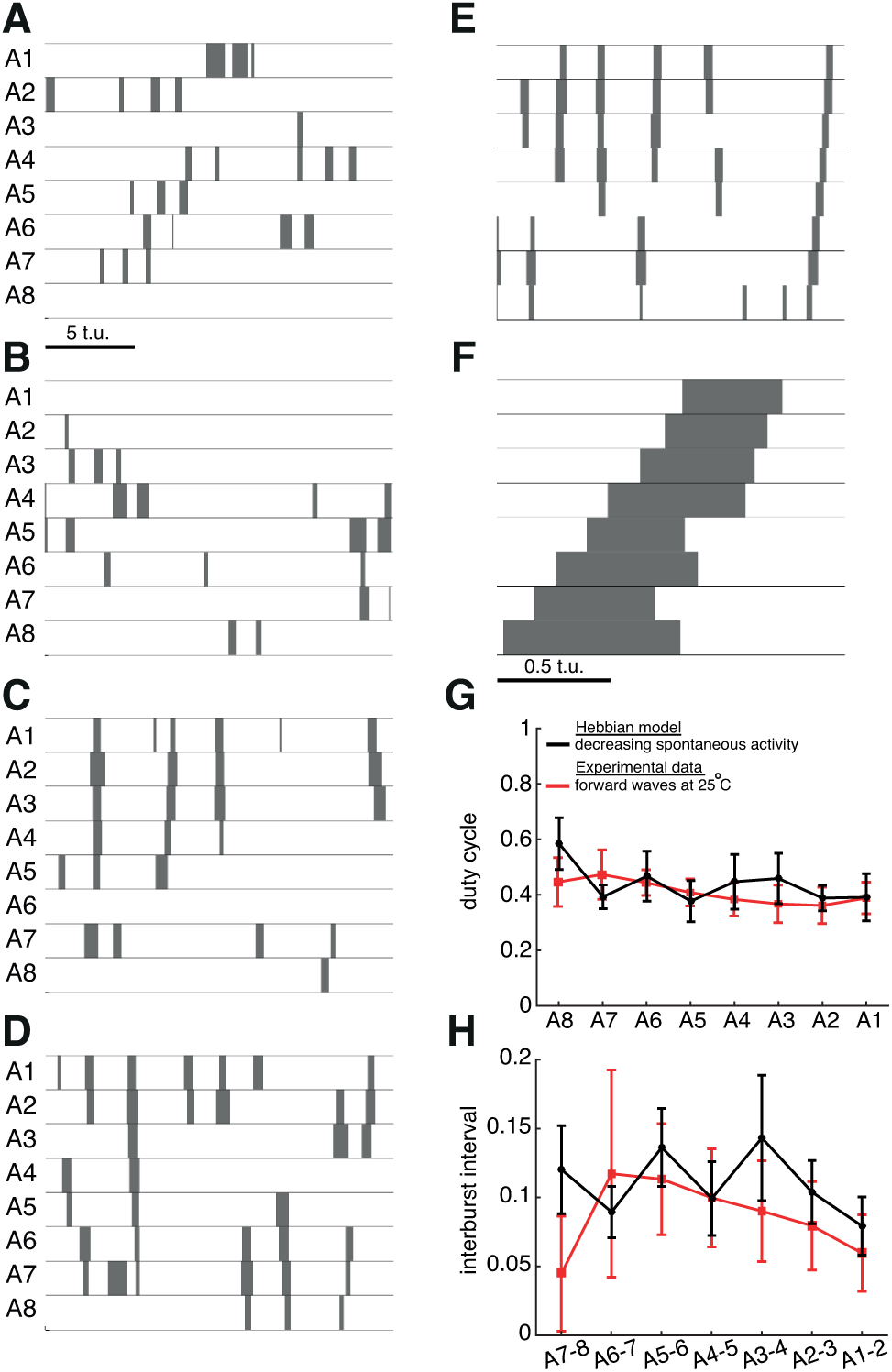
Emergence of coordinated output in the Hebbian efficacy model with additional end drive. **A-E**. Spatio-temporal patterns of supra-threshold excitatory activity at the developmental time denoted with arrows in Figure 4B. Scale bar in A applies to panels A to E. **F**. Applying *P*_ext_ = 1.7 to *E*_8_ in the network with the final set of weights generates a forward wave. **G, H**. Duty cycle and interburst interval (mean ± S.D.) for waves that were generated over 20 trials of wave development as in Figure 4C. The model is compared to experimental data as in Gjorgjieva et al. (2013).

The Hebbian efficacy model with balanced activity across the segments generated stable weight distributions even when the initial conditions for the weights were randomly distributed in a given range, albeit still weak (Figure 4C). Increasing that range of initial conditions fails to produce stable weights (data not shown). To determine if the final weights can indeed generate waves, forward waves were initiated by driving the most posterior population *E*_8_ in twenty networks where the weights were generated from random initial conditions (Figure 6F). The resulting waves show variable interburst intervals and variable duty cycles across the different segments that often fall outside the experimentally measured ranges (Figure 6G,H).

### Wave sensitivity to spontaneous activity patterns in the Hebbian efficacy model

How sensitive is weight development and wave generation in the model to changes in the properties of spontaneous activity? Modest changes to the distribution for the generation of spontaneous input strength *R_i_* preserve the interburst intervals, but retain the irregular duty cycles during wave propagation (Figure 7). Some cases, however, produce a stable configuration of weights too small to generate robust waves (data not shown). More substantial variations in the spontaneous input strength result in more significant degradation of the final weight distributions and ultimate failure of the network to generate waves. Similarly, stable weight distributions and propagating waves are produced only for a small subset of efficacy time constants recovery parameters. Mistuning these parameters either destabilizes the weights so that they never saturate to a stable value, or stabilizes the weights too early before they are sufficiently strong for wave generation (data not shown).

**Figure 7.**
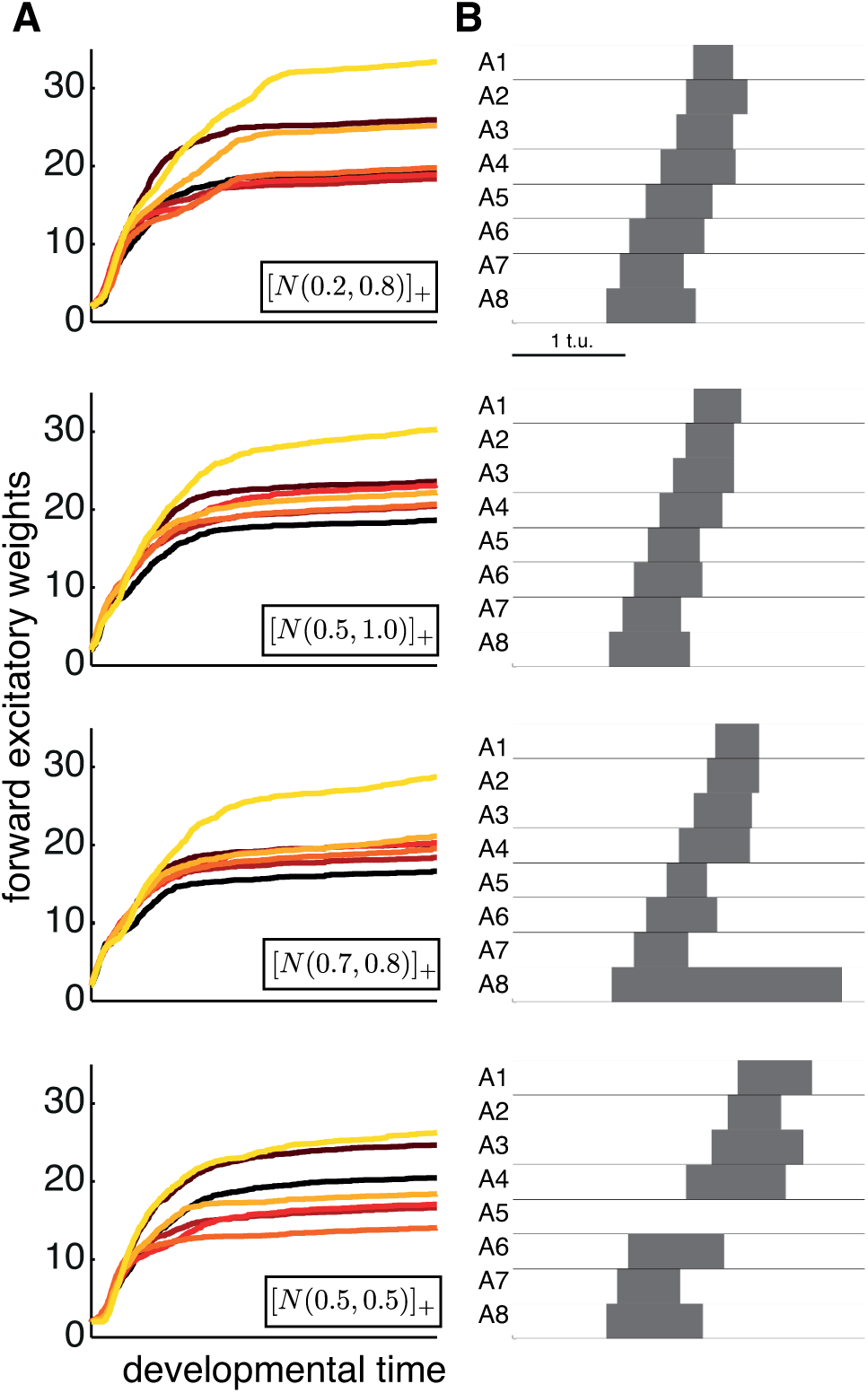
Varying the mean and variance of spontaneous input in the Hebbian efficacy model. **A**. Forward excitatory weights (as per Figure 3). The mean (*μ*) and the S.D. (*σ*) of the distribution for *R_i_* ~ [*N*(*μ*, *σ*)]_+_ were varied as shown in the inset of each plot. Every 8,000 time units, *σ* was decreased by 0.05. Compare to Figure 4 where *μ* = 0.5 and *σ* = 0.8. **B**. Examples of forward waves generated with the steady state weights at the end of each simulation in A.

### Conclusion for the Hebbian model(s)

For our networks with predetermined connectivity spanning only neighboring segments, we conclude that the basic requirement for coincident pre- and postsynaptic activity in the Hebbian model is insufficient to generate functional weight distributions that produce propagating waves with the appropriate timing relationships. Although model performance improved with the introduction of new assumptions (synaptic efficacy and additional spontaneous drive to the end segments), the final weight distributions and waves were highly variable. This suggests that Hebbian mechanisms likely play a minor role in tuning the motor network for wave propagation.

### The homeostatic model

When experimentally challenged with perturbations in synaptic structure and function, neurons have the remarkable ability to regulate their synaptic strengths back to the normal range. Various homeostatic mechanisms that maintain neuronal stability have been identified in invertebrates and vertebrates (Davis, 2006; Marder, 2012; Marder and Goaillard, 2006; Perez-Otano and Ehlers, 2005; Pozo and Goda, 2010; Turrigiano, 2008). One of the best-defined systems for analysis of the homeostatic mechanisms that regulate synaptic efficacy is the *Drosophila* neuromuscular junction (NMJ). The NMJ exhibits a strong homeostatic response to changes in postsynaptic excitability (Davis et al., 1998; Petersen et al., 1997). In contrast to the popular view that homeostatic plasticity is a slow phenomenon involving many neurons simultaneously, neurons may also undergo rapid synaptic tuning (Pozo and Goda, 2010).

Therefore, we considered a different model to regulate synaptic strength: a *homeostatic* mechanism that maintains network activity at stable levels based on spontaneous input. In our implementation of the homeostatic model, synaptic weights are potentiated or depressed to maintain a constant average activity of the excitatory population in each segment (see Methods). Spontaneous input is necessary for synaptic change. As such, our homeostatic model falls under the category of local activity-dependent models which require that each neuronal population computes a running average of its activity and the weights are modified to maintain that quantify (Zenke et al., 2013). A different class of homeostatic models are non-local models, which requires that all incoming weights to neuronal population are kept normalized even in the absence of activity (such as synaptic scaling) (Turrigiano et al., 1998).

The mechanisms for bidirectional homeostatic plasticity (simultaneously inducing potentiation and depression) have recently been demonstrated for the *Drosophila* NMJ (Gaviño et al., 2015), and also in *Drosophila* central neurons. The motor and visual system of developing larvae also show bidirectional structural homeostasis (Tripodi et al., 2008; Yuan et al., 2011), as well as the mushroom body of the adult (Kremer et al., 2010). In our homeostatic plasticity model, weights potentiate or depress to maintain excitatory activity in all segments (postsynaptic to both excitatory and inhibitory weights) at the same target level. For instance, weights potentiate or depress based on whether the average excitatory activity *r_i_* in a segment *i* is larger than the modification threshold, *θ*(*r_i_*); the threshold itself is a nonlinear function of the slow average of postsynaptic activity, as proposed in the BCM plasticity model (Bienenstock et al., 1982) (Figure 8A,B). To determine the steady state activity, we can solve for the case when the weights are also at a steady state. Then from equations (15)–(19)

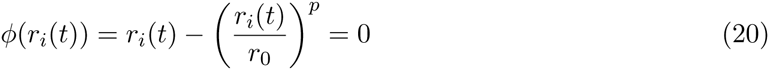
and the steady state excitatory activity is

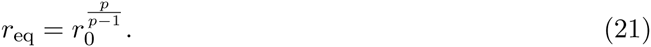

**Figure 8.**
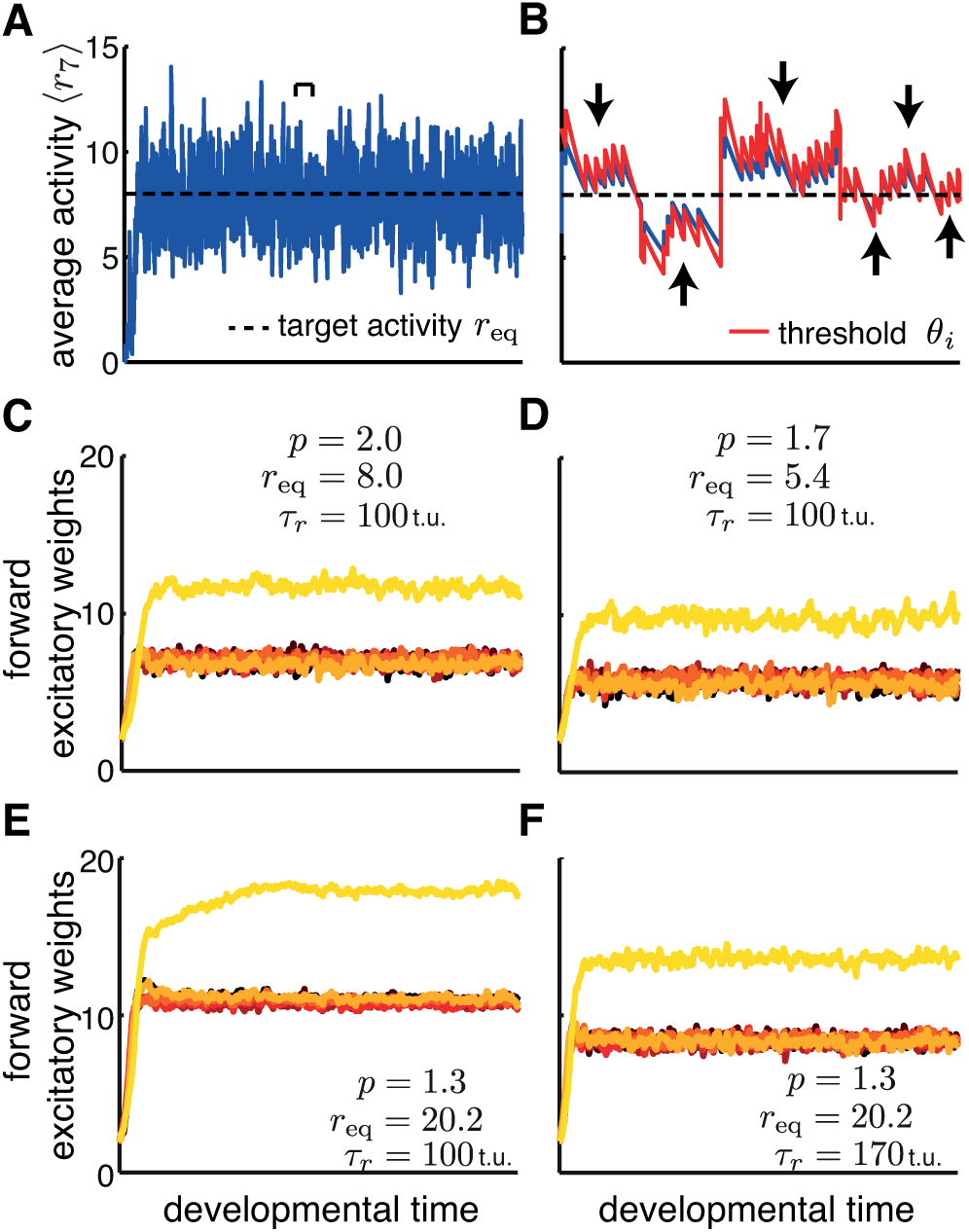
The average segmental excitatory activity and weights in the homeostatic plasticity model. **A**. The average excitatory activity of one segment, *E*_7_ is shown as a function of time of development fluctuating around the target activity. Spontaneous activity was generated with *R_i_* ~ [*N*(0.5, 0.4)]_+_ and *T_i_* ~ *U*(2, 3) throughout development. **B**. The modification threshold is a function of the average activity, and determines whether the weights will be potentiated (if the threshold is smaller the averaged activity) or depressed (if the threshold is above the averaged activity) as indicated by the upward and downward pointing arrows respectively. The threshold is shown for the time denoted by the bracket in A. **C,D**. Varying *p*, while keeping *τ_r_* constant, affects the target activity *r*_eq_ as in Eq. (21) and consequently the steady state feedforward excitatory weights. **E,F**. Varying *τ_r_* also affects the steady state weights, even when the target activity is the same. Here, spontaneous activity was generated with *R_i_* ~ [*N*(0.5, 0.4)]_+_ and *T_i_* ~ *U*(2, 3). Activity remained constant during development. In C-F, weights shown as per Figure 3, but only forward excitatory weights for illustration.

When *r*_0_ = 2.0 and *p* = 1.5, then *r*_eq_ = 8.0 (Figure 8A). Varying *p* affects the value of the steady state excitatory activity *r*_eq_ (Eq. 21) and the steady state weights in the network (Figure 8C). For instance, when *p* = 1.7, maintaining *r*_eq_ = 5.4 requires smaller weights in the network (Figure 8D). Similarly, when *p* = 1.3, maintaining *r*_eq_ = 20.2 requires larger weights (Figure 8E). The timescale *τ_r_* also has an effect on the steady state weights. Increasing *τ_r_* results in lower average excitatory activity, *r_i_*, (Eq. 13) and smaller steady state weights (Figure 8F). Therefore, the parameters in the homeostatic plasticity model determine the average activity of the excitatory populations and the strength of the steady state weights in the network necessary to maintain that activity.

### The homeostatic model produces stable weight distributions

Figure 8 shows that even at a constant level of spontaneous network activity during a simulation of development, the weights stabilize to maintain the target level of postsynaptic activity. If the amount of spontaneous activity decreases during the developmental period as the network assembles (Crisp et al., 2008), then the weights increase more gradually (Figure 9A), with a time constant combining the slow synaptic weight change and the decrease in spontaneous activity. Weight evolution goes through a plateau before the network has fully assembled. This plateau occurs as a result of the interaction of spontaneous input and recurrent activity in the network when the weights strengthen sufficiently to propagate activity to neighboring segment. Changing the properties of spontaneous activity can change when (and whether at all) this plateau occurs (Figure 11).

**Figure 9.**
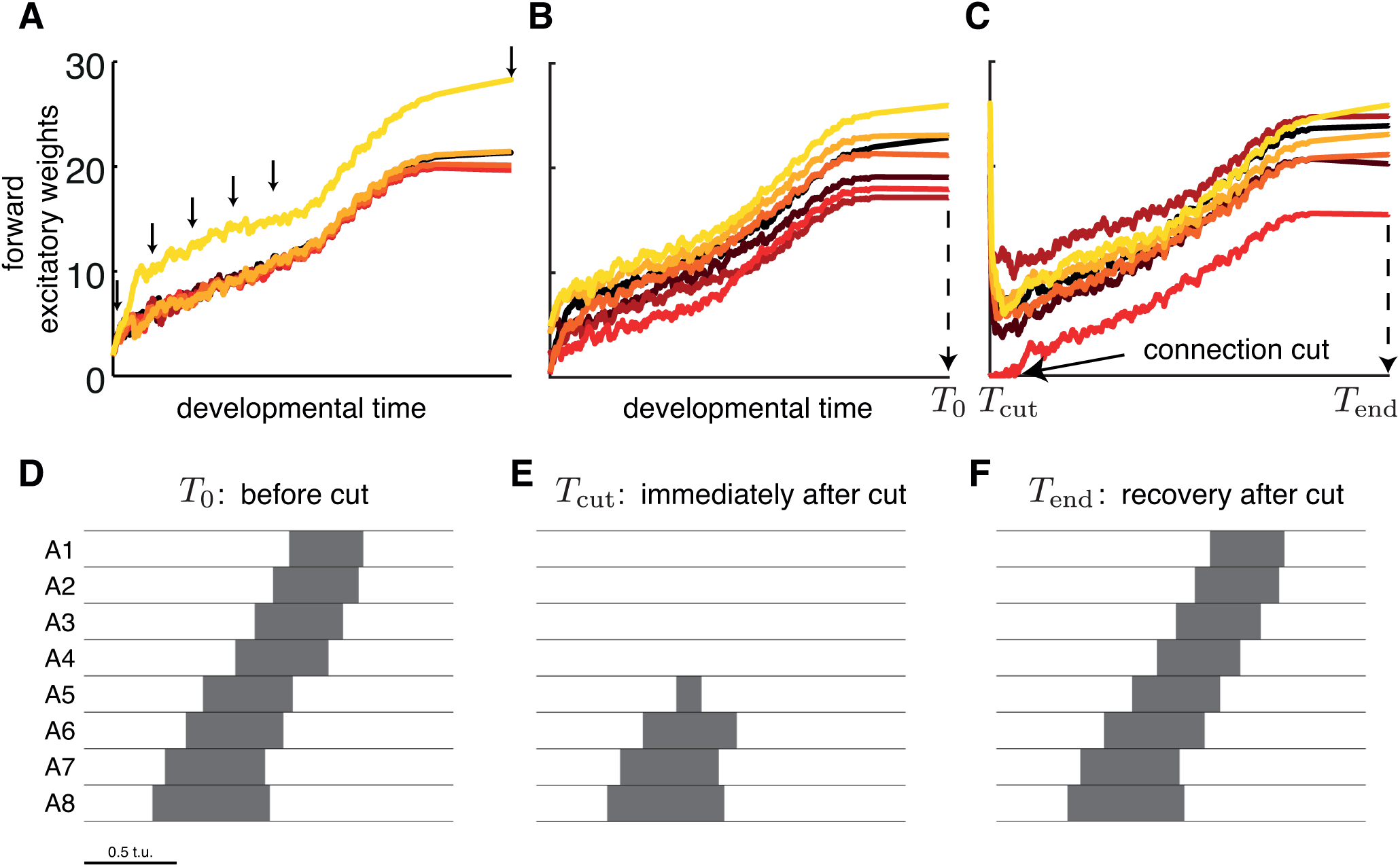
Weight development in the homeostatic model with decreasing spontaneous activity in development. **A**. The development of the weights under the homeostatic model (16)–(19) from an identical initial condition(*b* = −*d* = 2). Spatio-temporal patterns of excitatory activity at the developmental time denoted with arrows are shown in Figure 10. **B**. Same as A but with random initial conditions starting in the range of 0 and 5. **C**. Starting with the final weights in B (*T*_0_), only those between segments A4 and A3 were cut (weights set to 0) as indicated by the arrow (*T*_cut_), and allowed to develop again under the homeostatic model (*T*_end_). **D**. Forward wave generated using the final weights in B at *T*_0_. **E**. The network fails to generate waves when the connections are cut using the initial weights in C at *T*_cut_. **F**. Forward wave generated by the recovered network using the final weights in C at *T*_end_. In all cases spontaneous activity was generated with *R_i_* ~ [*N*(0.3,0.8)]_+_ and *T_i_* ~ *U*(2, 3). Every 8,000 time units, the standard deviation (*σ* = 0.8) of the distribution for *R_i_* was decreased by 0.04. Only excitatory weights in the forward direction are shown in A, B and *c* as per Figure 3.

During forward wave propagation, all but one of the excitatory weights 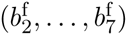 have as postsynaptic activity the excitatory activity of a segment in the middle of the body (*E*_2_,… *E*_7_). The postsynaptic activity for 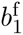 is *E*_1_, which is coupled only to a single neighboring segment, *E*_2_. Thus, to achieve the same target activity, 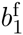 potentiates more strongly than the other forward excitatory weights (Figure 9A). The homeostatic model generates stable and functional weight distributions even when the weights start randomly distributed in a given range (Figure 9B,D). In contrast to the Hebbian model, the homeostatic model successfully generates stable weight distributions for larger ranges of initial conditions. For especially large ranges of initial weights, functional connectivity does not emerge as some weights become potentiated, while others depressed depending on their strength (data not shown).

Lastly, the homeostatic model can maintain weights at functional ranges even when network connectivity is perturbed. Following appropriate weight development in our functional model network, we ‘cut’ all connections between segments A3 and A4 (by setting the weights to 0) which eliminated wave generation (Figure 9E). Allowing the homeostatic model to act in the presence of spontaneous input recovered functional weights and appropriately timed waves (Figure 9C,F).

### The homeostatic model generates appropriately timed propagating waves

To examine the output of the model during development, we recorded the spatio-temporal patterns of activation in the network (Figure 10A–E) at the developmental time points denoted with arrows in Figure 9A. Coordinated output gradually improves in the network similar to the gradual improvement of motor output during *Drosophila* development (Crisp et al., 2008).

Are the stable weight configurations produced by this model able to generate robust propagating waves with appropriate timing relations? Probing twenty networks, where the weights were generated from random initial conditions, shows that the homeostatic model can indeed generate waves with regular interburst intervals and duty cycles across the different segments that closely match experimental variability (Figure 10G,H). This demonstrates that the homeostatic model successfully tunes network connectivity to a functional state which can generate propagating unidirectional waves with the appropriate segmental timing relations.

**Figure 10.**
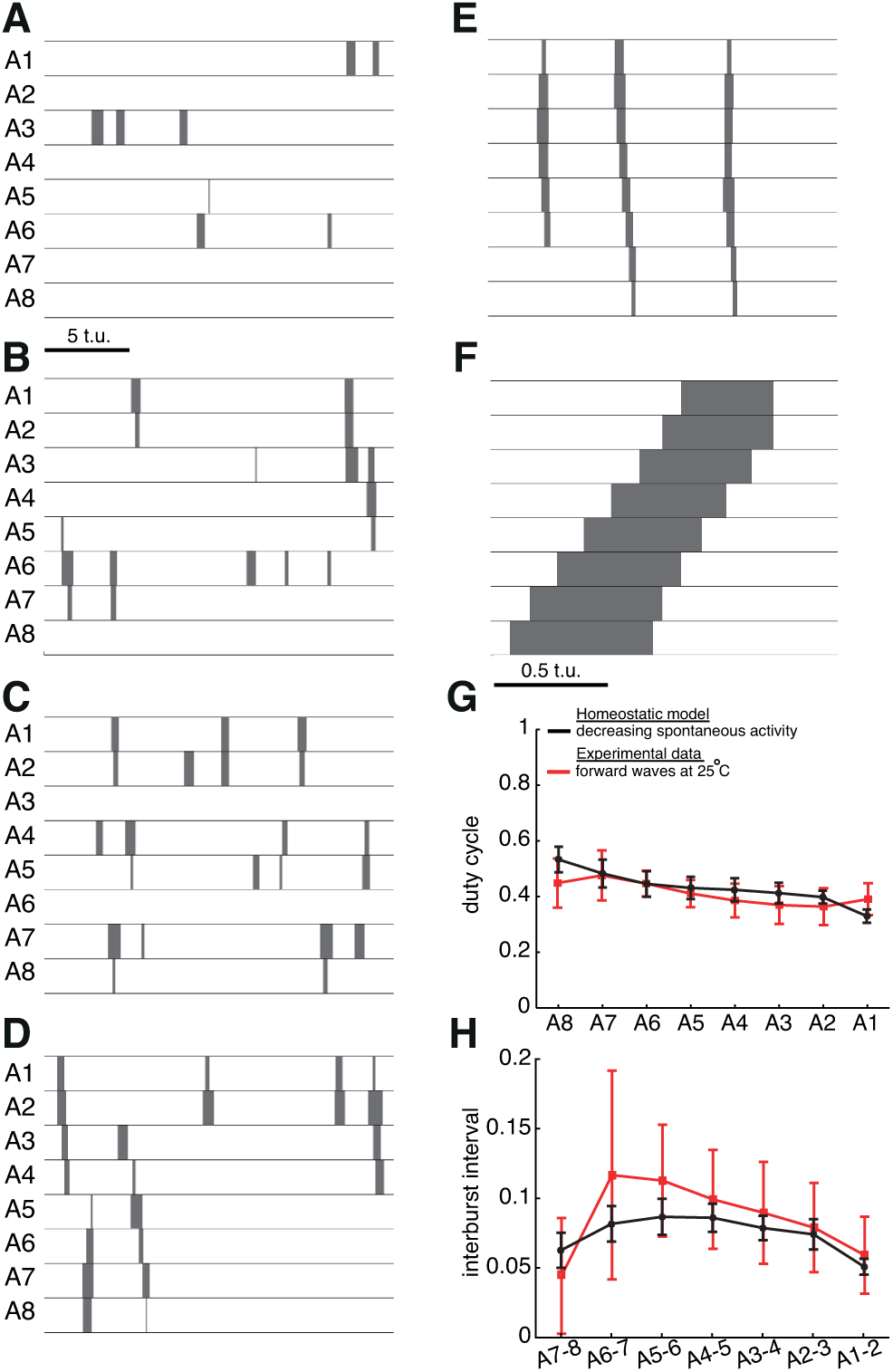
Emergence of coordinated output in the homeostatic model. **A-E**. Spatio-temporal patterns of supra-threshold excitatory activity at the developmental time denoted with arrows in Figure 9. Scale bar in A applies to panels A to E. **F**. Applying *P*_ext_ = 1.7 to *E*_8_ in the network with the final set of weights generates a forward wave. **G, H**. Duty cycle and interburst interval (mean ± S.D.) for waves that were generated over 20 trials of wave development as in Figure 4C. The model is compared to experimental data as in Gjorgjieva et al. (2013).

### Wave sensitivity to spontaneous activity patterns in the homeostatic model

The amount of spontaneous activity, determined by the properties of the distribution of input strength into each excitatory population, determines the steady state value at which the network weights saturate. To further investigate how the steady state weights depend on the nature of spontaneous activity, we examined weight development and wave generation as a function of the mean *μ* and the standard deviation σ of the distribution for spontaneous input strength *R_i_* ~ [*N* (*μ*, *σ*)]_+_.

Increasing *μ* raises the average spontaneous input strength, *R_i_*, thus increasing the excitatory activity in each segment, *E_i_*. Since *E_i_* receives two types of input (spontaneous and synaptic input), to maintain an average activity at the target level *r*_eq_, the synaptic input from neighboring populations must decrease; this can be achieved by decreasing the steady state strength of the weights (Figure 11). Waves generated by the network using the final steady state weights become longer as *μ* increases, and excitatory activity remains above threshold for a shorter time in each segment, resulting in shorter duty cycles and longer interburst intervals. Increasing *μ* ≥ 0.5 makes the influence of spontaneous input on excitatory activity so strong that very little drive from neighboring segments is needed to maintain the average excitatory activity at target. Therefore, the weights become too small for the network to generate waves (data not shown).

**Figure 11.**
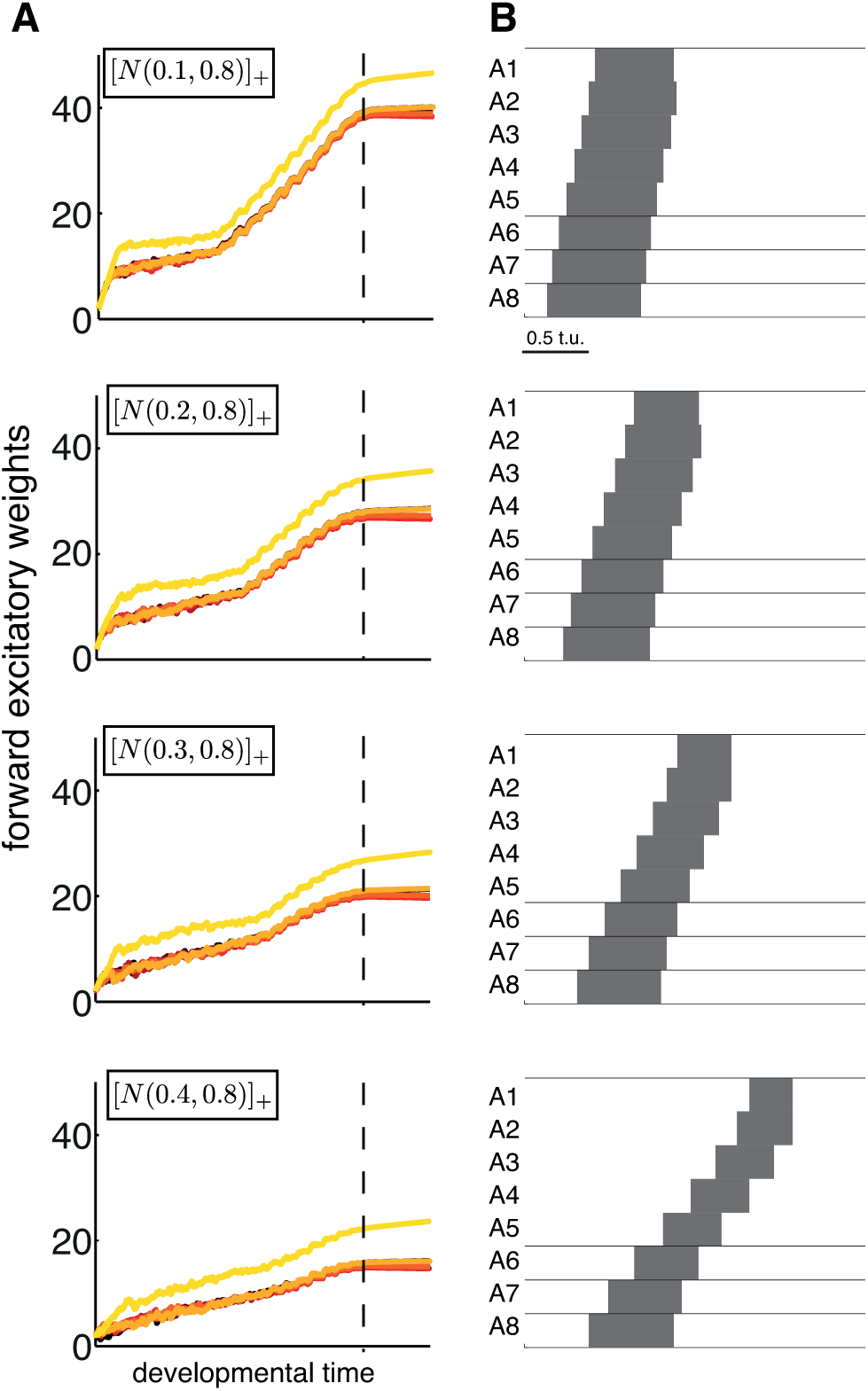
Varying the mean of spontaneous input in the homeostatic plasticity model. **A**. Forward excitatory weights. The mean (*μ*) of the distribution for *R_i_* ~ [*N*(*μ*, *σ*)]_+_ was varied as shown in top-left of each graph. In all cases *σ* = 0.8 and decreased over development by Δ*σ* = 0.04 every 8,000 time units. The time when the standard deviation reached 0 is denoted with vertical dashed lines. Weights shown as per Figure 3, but only forward excitatory weights for illustration. **B**. Forward waves generated with the steady state weights at the end of each simulation in A.

This suggests that for the homeostatic model to produce stable and functional network weights spontaneous activity must operate within a range. It should be sufficiently frequent to drive excitatory activity above threshold and to enable weight potentiation; yet it should not be too frequent, to allow steady-state population activity to be maintained by synaptic input from neighboring populations. The model is robust to changes in the amount by which spontaneous activity decreases during development and it can also generate stable and functional weights even when spontaneous activity is constant during development (Figures 8 and 10G,H).

### Conclusion for the homeostatic model

These results demonstrate that based on the properties of spontaneous activity, different stable weight configurations can be achieved with the homeostatic model by maintaining a target level of postsynaptic activity for each weight. The weights can be reliably reproduced over multiple simulation runs and initial conditions, and no additional assumptions are necessary to bound synaptic growth as for the Hebbian model. A network with the final set of weights can generate propagating waves with precise timing properties, such as regular interburst intervals and duty cycles across the segments. This suggests that differences in the nature of spontaneous activity during development may underlie variability of network connectivity and the resulting wave properties produced by the final network. Therefore, developmental differences in spontaneous activity (endogenous or environmental) represent one plausible way by which such variability is achieved in the motor network of *Drosophila* (Berni et al., 2012; Gjorgjieva et al., 2013).

## Discussion

How are neural circuits organized to maintain stable function and produce robust output? This task is made especially difficult during circuit development, when the properties of different circuit components are immature and changing, and input from the environment unreliable. Here we studied a model network of recurrently connected excitatory and inhibitory neuronal populations segmentally repeated in a single dimension, motivated by the motor network for *Drosophila* larval crawling. A key feature of this network, when appropriately tuned, is its ability to generate robust forward and backward waves of activity by driving each end of the network (Gjorgjieva et al., 2013). Examining sets of parameters that enable wave generation in the network has revealed nontrivial solution regions. We sought to identify plausible mechanisms by which these parameters emerge.

### Activity-dependent tuning of network connectivity

While there is ample evidence that spontaneous activity patterns instruct refinement of circuit connectivity in sensory systems (Huberman et al., 2008), the importance of spontaneous activity for the development of motor circuits is conflicting (Li et al., 2008; Marder and Rehm, 2005; Myers et al., 2005; Roberts et al., 2014). Network models of spontaneous activity in developing motor circuits have typically addressed the generation and properties of high-activity episodes interrupted by quiet periods in the embryonic chick spinal cord (Marchetti et al., 2005; Tabak et al., 2010, 2001, 2000). However, unlike sensory systems, it is unclear if this activity helps refine developing motor circuits.

Imaging of spontaneous activity during embryogenesis in *Drosophila* larvae has revealed a gradual progression of motor output, and manipulations of this activity point to the role of activity-dependent mechanisms in the tuning of the network (Crisp et al., 2011, 2008; Giachello and Baines, 2015). Using computational modeling we have demonstrated that spontaneous activity can indeed tune a weakly connected recurrent network if the appropriate activity-dependent tuning rules are used. We examined two styles of activity-dependent mechanisms for weight development, that through gradual improvement in network output lead to stable weight distributions to generate propagating waves with regular interburst intervals and duty cycles.

Hebbian-style models modify synaptic strength based on coincident pre- and postsynaptic activity. While Hebbian mechanisms instruct activity-driven refinements in the developing visual system (Huberman et al., 2008), applying similar mechanisms for the tuning of a motor network for wave propagation yields less successful results. Unlike sensory systems where neurons topographically project from an input to a target layer, the motor network in *Drosophila* larvae is segmentally organized and recurrently connected. Although correlated activity emerges between neighboring segments even when spontaneous input is uncorrelated, there is no obvious notion of pre- and post-synaptic: the same segment can be both presynaptic and postsynaptic to different weights. Thus, Hebbian mechanisms are less likely to play a fundamental role in fine-tuning network connectivity. Indeed, both Hebbian (bidirectional and efficacy) models fail to produce functional bidirectional weights that generates waves with appropriate timing relationships.

In systems as diverse as the crustacean stomatogastric system and the vertebrate visual system, homeostatic mechanisms control neuronal function through the regulation of synaptic efficacy and the modulation of intrinsic ionic conductances (Davis, 2006; Marder, 2012; Marder and Goaillard, 2006; Perez-Otano and Ehlers, 2005; Pozo and Goda, 2010; Turrigiano, 2008). Several forms of homeostatic plasticity have been proposed to stabilize network activity, such as regulation of the strength of synaptic transmission and synaptic scaling as a function of global network activity (Gonzales-Islas and Wenner, 2006; Turrigiano, 1999; Turrigiano et al., 1998; Turrigiano and Nelson, 2004). Altered activity patterns can scale synaptic connections through homeostatic mechanisms also in motor circuits (Borodinsky et al., 2004). Our homeostatic model was based on the goal to maintain a target level of excitatory postsynaptic activity, which was achieved by modifying the synaptic weights for which the target population is postsynaptic. Spontaneous activity was necessary to drive weight modifications, thus our model differs from global mechanisms like synaptic scaling (Turrigiano et al., 1998) and is more similar to the local BCM model (Bienenstock et al., 1982; Zenke et al., 2013).

Without any additional assumptions, the homeostatic model achieves stable and functional bidirectional weight distributions that generate waves with regular duty cycles and interburst intervals. Moreover, different wave properties can be produced from different spontaneous activity patterns, providing a way to generate wave variability in the experimental system. Therefore, our results suggest that activity-dependent homeostatic mechanisms are more likely to tune weak bidirectional connectivity in a recurrent network than Hebbian mechanisms. During tuning, the network gradually improves the coordination of its output, matching the gradual emergence of coordinated output in developing *Drosophila* larvae. The resulting network can produce appropriately timed network-wide waves of activity that propagate in either direction despite bidirectional connectivity. This property can be matched to the ability of *Drosophila* larvae to crawl forward and backward with similar properties (Gjorgjieva et al., 2013). The homeostatic model can also restore functional weights following perturbations of connectivity in the presence of spontaneous input with similar patterns as used for developmental tuning. Thus, if activity-dependent homeostasis continues to operate after development to keep patterned activity robust, then the network must continue to receive external input. This is likely since the *Drosophila* larval motor network receives descending input during continuous crawling, as well as additional proprioceptive input from the environment.

### Limitations of the plasticity models

Because no activity-dependent plasticity rules have yet been identified in developing motor circuits, it was natural to assume here that inhibitory plasticity is similar to excitatory plasticity (notably, homeostatic adjustment of both excitatory and inhibitory connections occurs (Gonzales-Islas and Wenner, 2006)). In both activity-dependent models, excitatory and inhibitory weights developed in a balanced fashion, enabling the generation of waves propagating in either direction. Therefore, neither type of weight is stronger than the other at any time. This is currently at odds with experimental data, where at least initially, inhibition is not required to generate output patterns (Crisp et al., 2008).

In addition to regulation of synaptic connectivity, which is realized by both our Hebbian and homeostatic models, homeostatic regulation may occur at the level of intrinsic neuronal excitability through the modulation of intrinsic ionic conductances. Theoretical modulation rules have been successfully applied to the regulation of motor output in the stomatogastric nervous system of crustaceans (LeMasson et al., 1993; Liu et al., 1998; Marder and Goaillard, 2006; O’Leary et al., 2014). Our current rate-based framework precludes such studies based on conductance-based models, although it can incorporate changes in intrinsic excitability by modulation of the activation functions in the population model equations.

Mapping of circuit connectivity in EM volumes has revealed the existence of long-range connectivity that spans non-neighboring segments in the *Drosophila* motor network (personal communication with Albert Cardona). However, how long-range connectivity changes during embryonic development is still unknown. Thus, we based our models on a ‘minimal’ network architecture where the nature of connectivity (but not strengths) was predefined to be nearest neighbor (Gjorgjieva et al., 2013). As we add other types of long-range connections, we will need other kinds of mechanisms, some of them Hebbian, in combination with homeostatic plasticity (Vitureira and Goda, 2013; Zenke et al., 2013) to ensure that connectivity, short and long-range, is refined in the appropriate manner.

### Generality of the network model

Although our model is motivated by the production of motor output during crawling in *Drosophila* larvae, it is not an anatomical model of the *Drosophila* motor network and does not incorporate any detail about the organization and neural identity of different network elements such as interneurons, motor neurons and muscles; instead the activity of each segment is represented with a single excitatory and inhibitory population. Therefore, our model does not capture the known molecular underpinnings of the homeostatic mechanisms that regulate synaptic efficacy and channel function in *Drosophila* (Davis and Müller, 2015; Tripodi et al., 2008). For instance, what kind of a ‘sensor’ monitors neuronal or muscle activity, and by what mechanisms is pre- and postsynaptic function modulated?

Yet, the generality of the model allows it to be applied to other circuits where activity-dependent refinement of connectivity leads to appropriate connectivity for generating unidirectional propagation of activity. One example is the generation of spontaneous waves in the developing mammalian cortex; waves are produced by recurrent networks without a particular directionality of connections, but travel along stereotypical directions (Conhaim et al., 2010; Lischalk et al., 2009).

We conclude that for the fixed network architecture with nearest-neighbor connections, Hebbian mechanisms are unlikely to play a major role in activity-dependent tuning of connectivity in recurrent bidirectionally connected networks for wave propagation. By contrast, homeostatic plasticity mechanisms, which maintain target levels of activity using spontaneous input, succeed in generating reproducible patterns of network connectivity. As such, they are more likely than Hebbian-style mechanisms in regulating activity-dependent tuning of bidirectional motor networks for activity propagation. Both mechanisms are likely to be necessary for the refinement of networks with long-range connectivity.

## Acknowledgement

Funding: Cambridge Overseas Research Fund, Trinity College and Swartz Foundation (JG), Wellcome Trust VIP funding (JFE). JG is supported by a Career Award at the Scientific Interface from the Burroughs-Wellcome Fund. We would like to thank Sarah Crisp, Michael Bate and Matthias Landgraf for useful discussions, and members of the Marder Lab at Brandeis University for feedback on figures.

